# Mitigation of coral bleaching by antioxidants

**DOI:** 10.1101/281014

**Authors:** Michael Marty-Rivera, Loretta M. Roberson, Guillermo A. Yudowski

## Abstract

Coral bleaching, loss of symbiotic dinoflagellate algae from the coral holobiont, is a complex phenomenon that can result in coral death and reef degradation. Reactive oxygen species (ROS) have been suggested as a possible mechanism underlying this event. To determine if antioxidants can be used to reduce ROS production and coral bleaching, we tested the effects of thermal stress in *Aiptasia pallida* a model system for coral bleaching studies, and the scleractinian coral, *Porites astreoides.* We analyzed host ROS levels, symbiont dark-adapted quantum yield of photosystem II, and symbiont loss in the presence or absence of antioxidants. We found that a single dose of the antioxidant catechin, significantly reduced ROS levels in the hosts, mitigated the degradation of the symbiont’s quantum yield and reduced the loss of symbionts from thermally stressed *P. astreoides* but not from *A. pallida*. Taken together, these results support a key role of ROS and that antioxidants can prevent symbiont degradation and loss during thermally-induced bleaching in *P. astreoides.*

**Abbreviations**
PAM: Pulse-Amplified Modulation
ROS: Reactive Oxygen Species

## Introduction

Coral reefs have been degrading at an alarming rate over the past three decades (Gardner *et al.*, 2003; Eakin *et al.*, 2010). One of the major driving forces behind coral cover decrease is believed to be increases in sea surface temperature, where either an increase of 1 – 2 °C for several weeks, or 3 – 4 °C for a few days above the summer mean temperatures can result in coral bleaching (Hoegh-Guldberg and Smith, 1989; Glynn and D’Croz, 1990; Jokiel and Coles, 1990). Coral bleaching is an event where coral symbionts, *Symbiodinium* sp., lose photosynthetic efficiency or decline in density from the coral host tissue, resulting in a decrease in carbon and energy fluxes, and ultimately coral death (Fitt *et al.*, 2001; Weis, 2008). However, the underlying cellular mechanisms are less understood.

Excess reactive oxygen species (ROS) production has been proposed as a major mechanism leading to loss or degradation of symbionts from their coral host cells during the late phases of coral bleaching (Smith *et al.*, 2005; Lesser, 2006). ROS consist mainly of hydrogen peroxide (H_2_O_2_), superoxide anion radical (O_2_^.-^), and singlet state oxygen (^1^O_2_), which can cause damage to membranes, DNA, and protein denaturation of both symbiont and host cells leading to coral bleaching, and in some cases death (Lesser, 2006). ROS scavenging in cells is accomplished mainly by superoxide dismutase and ascorbate peroxidase, but the increase in ROS associated with thermal stress can exceed detoxification rates, resulting in deterioration and degradation of the holobiont (Matta and Trench, 1991; Smith *et al.*, 2005). Reductions in the maximum dark-adapted quantum yield of photosystem II from the symbiont (F_v_/F_m_), a measure of their photosynthetic efficiency, is observed in the early phases of natural bleaching events (Gates *et al.*, 1992; Franklin *et al.*, 2004), while a lethal ROS diffusion into the host cell is thought to occur in the terminal phase of the process (Weis, 2008). Previous studies have investigated coral’s antioxidant response, a measure thought to reduce thermally induced stress (Lesser, 1996; Yakovleva *et al.*, 2004; Krueger *et al.*, 2015). Here, we sought to test the hypothesis that increases in ROS induced by thermal stress are sufficient to trigger a reduction in photosynthetic efficiency and *Symbiodinium* sp. expulsion in the model system, *Aiptasia pallida* and the coral *Porites astreoides* (Green *et al.*, 2008; Weis *et al.*, 2008) and test the hypothesis that a reduction in ROS buildup by a naturally occurring antioxidant can prevent coral bleaching.

## Materials and methods

### Animal collection and maintenance

*A. pallida*, were purchased from Carolina^^®^^ Biological Supply Company (Burlington, NC, USA). Sixteen colonies of *P. astreoides* 3-5 cm in diameter were collected under the Puerto Rico Department of Natural and Environmental Resources permit No. 2016-IC-187 from Escambron Beach in San Juan, Puerto Rico (18°28’03”N 66°05’27”W) and transported to the lab in a cooler with ambient seaweater. Upon arrival, animals were transferred to holding tanks with artificial sea water (ASW) for *A. pallida*, or filtered sea water (FSW) for *P. astreoides*, and gradually acclimatized for 3 hours. Sea water was filtered using Whatman^^®^^ GF/F filters (Pore size: 0.7 µm) to ensure no free swimming *Symbiodinium* sp. entered our tanks. The selection of ASW, FSW and maintenance conditions were based on the animal’s place of origin. Sea anemones were maintained in 30 L tanks at 25-27 °C, 35-37 PSU, and an irradiance of 80-100 μmol quanta m^−2^ s^−1^ on a 12 h:12 h light:dark photoperiod. Corals were maintained in 9.5 L tanks at 27-29 °C, 35-37 PSU, similar to ambient conditions at the collection site at a depth of 2.0 meters, and light levels of 70-100 μmol quanta m^−2^ s^−1^ on a 12 h:12 h light:dark photoperiod. We selected this light level to have thermal stress as the principal variable impacting ROS production (Hawkins and Davy, 2013; Gibbin *et al.*, 2014; Matthews *et al.*, 2016). Both species were maintained in tanks with air and water pumps for at least a week and fed with freshly hatched brine shrimp, water was changed a day before feeding, and upon once upon experimental start. All animals were starved for 2-3 days prior to being exposed to experimental conditions.

### Experimental treatments and symbiosome isolation

Both anemones and corals were subjected to thermal stress and treated with a single 5 µM dose (final concentration) of the antioxidant (+)-catechin (Sigma-Aldrich, St. Louis, MO), dissolved in 70% ethanol administered at the start of the experimental period with no subsequent water changes in the tank for 4 days. We chose catechin due to its availability and extensive characterization (Mendoza-Wilson and Glossman-Mitnik, 2006). Catechin can be found in many plant species, especially fruits and green tea, traits attributed to catechin are that of being a ROS scavenger, and stress regulator (Chobot *et al.*, 2009). We used a relatively low antioxidant concentration of 5 µM compared to several studies in other marine organisms which have used up to 125 µM (Dunlap and Yamamoto, 1995; Lesser, 1997). High concentrations of antioxidants can have pro-oxidant and harmful effects (Villanueva and Kross, 2012; Sayin *et al.*, 2014).

#### Aiptasia pallida

A total of 28 *A. pallida* (2-3 mm oral disc diameter) were randomly selected and acclimated to 28-29 °C for 24 hours before transferring to experimental tanks. Pre-acclimated anemones were then transferred into 0.5 L tanks in the presence or absence of 5 μM catechin, then kept for four days at 33-34 °C, 35-38 PSU, and irradiance of 85-95 μmol quanta m^−2^ s^−1^ on a 12 h:12 h light:dark photoperiod. Prior studies have shown that temperatures > 33 °C induce bleaching in *A. pallida* (Paxton *et al.*, 2013). Control experiments were carried out identically but maintained at the baseline temperature of 25-27 °C.

#### Porites astreoides

Sixteen corals were randomly selected from the holding tank and transferred into a pre-acclimation tank at 29 °C for 24 h to avoid thermal shock from a rapid temperature change; four corals were used in each treatment, as independent replicates. After 24 h, they were transferred to 9.5 L tanks with either 5 μM of antioxidant or FSW. They were kept for four days at 32-33 °C, 35-37 PSU, and irradiance of 70-90 μmol quanta m^−2^ s^−1^ on a 12 h:12 h light:dark photoperiod. Control experiments were carried out identically but maintained at the baseline temperature of 27-29 °C.

After treatments, tissues were harvested for symbiosome (*Symbiodinium* sp. encased in a host membrane) isolation at the end of a dark-phase. Whole *A. pallida* individuals (mean wet weight 0.28 g) were scraped from the tank using a blunt edged tool, weighed, and then homogenized in a set volume of 5 mL ASW using a Teflon-glass tissue grinder. The tissue slurry was then centrifuged at 3200 *g* for 10 min to remove tissue supernatant from the symbiosomes and the pellet was resuspended in 10 mL ASW. This homogenized symbiont slurry was used for microscopy analysis. *P. astreoides* symbiosomes were treated in the same manner except tissue was stripped from the entire coral skeleton of each colony using a water jet with a constant 100 mL of FSW (Interplak^®^, CONAIR^®^) prior to centrifugation.

### Microscopic analysis

#### Reactive oxygen species

ROS levels were investigated in freshly isolated host cells still harboring their symbionts using a ROS sensitive dye and analyzed using confocal microscopy at the end of a dark-phase. Single host cells along with their symbionts (symbiosomes), were identified in an equatorial plane by their characteristic plasma membrane surrounding individual *Symbiodinium* sp. which autofluoresce at 650 - 725 nm due to their chlorophyll content (Fig. 1A and B).

**Fig. 1.**
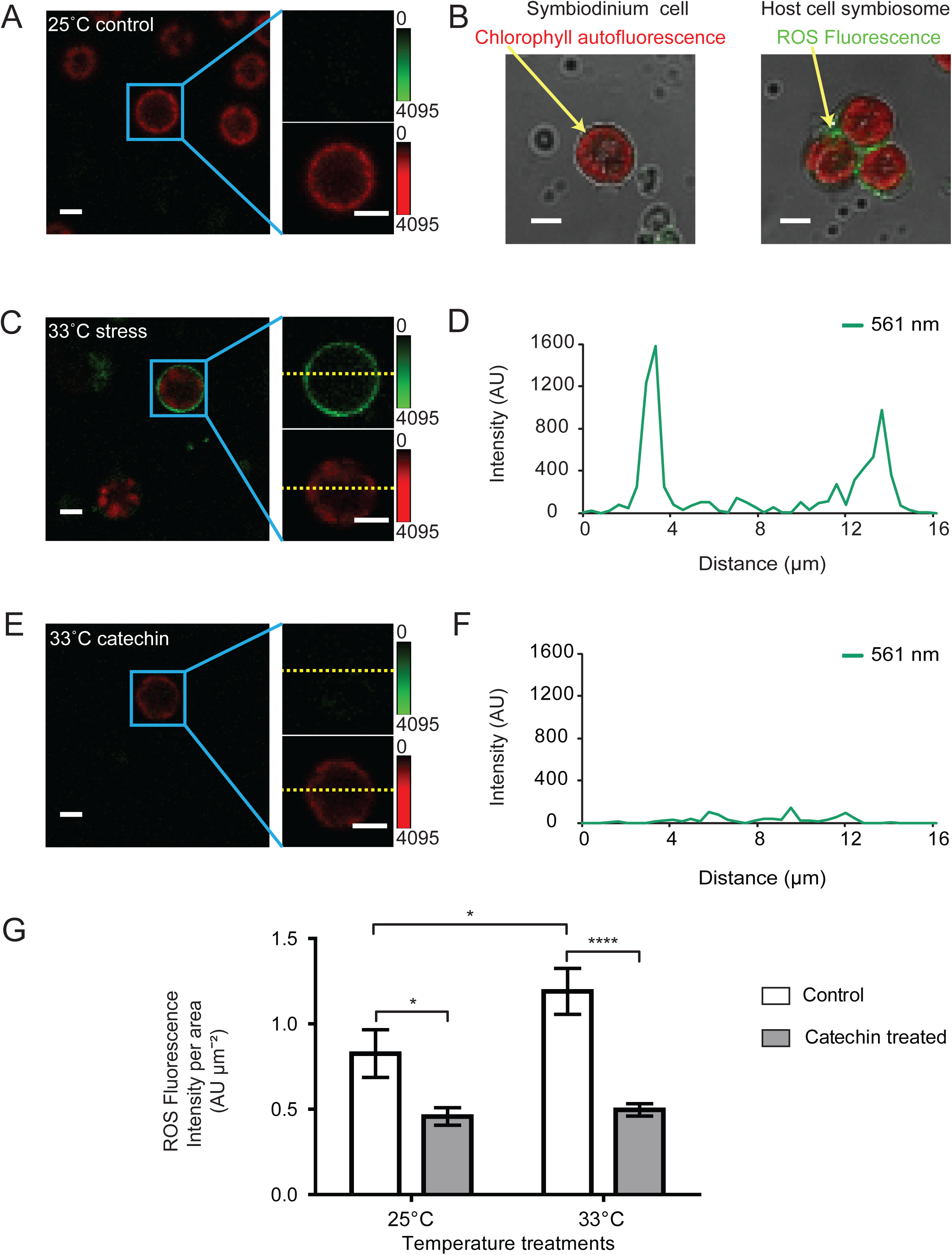
ROS in *A. pallida* cell symbiosomes. (A) Isolated control symbiosomes imaged under confocal microscopy in the presence of a ROS sensitive dye presented undetectable signal (green channel insert, top) and characteristic chlorophyll autofluorescence (insert, bottom). Colored gradient bar represents fluorescence min/max values. (B) Isolated symbiosomes from the sea anemone *A. pallida* were readily identified by their host cell membranes containing autofluorescent symbionts, ROS fluorescence is also depicted in green. (C) Symbionts from thermal stress treatment presented significant fluorescence from the ROS sensitive dye (green) (D) Line intensity profile of ROS fluorescence in stress treated symbiosomes (from yellow line in top insert of Fig. 1C). (E) Catechin + thermal stress treated symbiosomes incubated with CellROX ^®^. (F) Line intensity profile of the catechin + thermal stress treated symbiosome from E. (G) Catechin significantly reduced ROS fluorescence in control and thermally stressed symbionts after a 4-day treatments (2-Way ANOVA: temperature factor *P* = 0.0490; antioxidant factor *P* < 0.0001; temperature x antioxidant factor *P* = 0.1120). Error bars represent the SE. *n* = 4 per treatment; sample size per *A. pallida* = 75. Asterisks denote statistical significance of the post-hoc analyses. Scale bar = 5 μm.

ROS was measured in extracted symbiosomes by confocal microscopy using the nonspecific ROS detection probe CellROX^®^ Orange Reagent (Life Technologies, Eugene, OR, USA) at excitation wavelength 545 nm and emission 560 - 590 nm. Chlorophyll autofluorescence in both anemones and corals was measured using excitation wavelength of 488 nm and emission range of 650-750 nm to avoid signal interference with the CellROX probe (Fig. S1). For staining, 0.49 ml of homogenized symbiont slurry was incubated with CellROX^®^ Orange in ASW or FSW to a final 1 ml solution of 5 μM for 30 min at 32.5 °C in darkness. After incubation, samples were washed with ASW or FSW and imaged by confocal microscopy. Imaging was performed using a Nikon A1+ with a 60x/1.40 0.21 mm working distance oil immersion objective (Nikon Instruments Inc., NY, USA). Images were analyzed using the open source ImageJ/Fiji software (Schindelin *et al.*, 2012). Single symbiont cells were randomly selected for analysis. ROS was measured by excitation intensity in whole symbiosomes selected as Regions of Interest (ROI). Host cell membranes were identified using Differential Interference Contrast (DIC) microscopy (Fig.1B). Line scan selection measurements of ROS fluorescence intensities were done to determine the distribution of fluorescence in the symbiosomes using the line selection tool along with plot profile.

#### Photosynthetic efficiency

To investigate if antioxidant treatment could prevent the impairment of the photosynthetic function observed under thermal stress, we measured maximum quantum yield of photosystem II (F_v_/F_m_) at the end of a dark-phase in freshly isolated symbionts by microscopy-based Pulse-Amplitude Modulated fluorometry (PAM). A total of 50 μl of the homogenized symbiont slurry per individual was used for analysis. A Microscopy-PAM (Zeiss Axio Scope.A1 (Zeiss, Göttingen) and IMAG-CM; Heinz Walz GmbH, Effeltrich, Germany) was used to measure F_v_/F_m_ in randomly selected individual cells as indicated in figure legends, where variable fluorescence (F_v_) was calculated by subtracting basal fluorescence (F_o_) from maximum fluorescence (F_m_) in dark-adapted samples (Genty *et al.*, 1989). Fluorescence excitation was provided by a saturating pulse of a 470-nm light module (IMAG-L470M).

#### Symbiodinium sp. density

Isolated *A. pallida* and *P. astreoides* symbionts were counted using an hemocytometer. A total of 40 μl of symbiont slurry was used in the analysis. four center squares counts were made for each treatment using 10 μl. We calculated *A. pallida* cell density per gram by normalizing our cell counts per area to total sea anemone wet weight, while *P. astreoides* counts were normalized by chamber volume.

#### Spectral analysis

Emission wavelength for *P. astreoides* symbionts was measured with a Nikon A1+ spectral configuration detector from 421-741 nm at a resolution of 10 nm to ensure there was no overlap between CellROX^®^ Orange and symbiont autofluorescence. 50 μl of the symbiont slurry was used to look at 4 individual *P. astreoides* symbionts.

### Statistical analysis

All data for both anemones and coral (ROS levels, photosystem II photochemical efficiency and symbiont cell counts) were analyzed by 2-way ANOVA with temperature and antioxidant treatment as factors and a Bonferroni multiple comparison post-hoc test. Significant statistical difference was set to α = 0.05. Data analysis was carried out using GraphPad Prism version 6.01 (San Diego, CA, USA).

## Results

### Natural antioxidant and ROS in *A. pallida*

ROS levels were investigated in freshly isolated *A. pallida* symbionts using a ROS sensitive dye and analyzed using confocal microscopy. Single host cells along with their symbionts (symbiosomes), were identified by their characteristic plasma membrane surrounding individual *Symbiodinium* sp. which autofluoresce at 650 - 725 nm due to their chlorophyll content (Fig. 1A and B). *A. pallida* controls exhibited an average intensity of 0.826±0.140 AU μm^−2^ (Mean±SE; Figs. 1A and G). These levels could indicate basal physiological levels of ROS or low levels of stress. Individual cells from thermally stressed anemones had a significantly higher ROS signal of 1.190±0.135 AU μm^−2^ (Figs. 1C and G). A representative line scan from individual cells shows increased ROS levels in the host cellsurrounding *Symbiodinium* sp. (Fig. 1D). Thermally stressed anemones incubated with catechin and control anemones in the presence of catechin yielded a lower signal of 0.496±0.036 and 0.457±0.051 AU μm^−2^, respectively (Figs. 1E, F and G), suggesting a decrease in ROS fluorescence by the incubation with 5 µM catechin (Fig. 1 G). Additional statistical information is included in supplemental materials (Table S1).

### Dark-adapted maximum quantum yield of photosystem II of *A. pallida*

Our microscopic PAM analysis of single symbionts indicated that F_v_/F_m_ values (Mean±SE) from control experiments were 0.558±0.004, similar to those previously described in the literature for *Symbiodinium bermudense*, which can inhabit *A. pallida* (Figs. 2A and D) (Lesser, 1996; Perez *et al.*, 2001). Thermally stressed anemones had a similar F_v_/F_m_ mean value of 0.545±0.012 (Figs. 2B and D). While stressed anemones incubated in the presence of catechin yielded higher F_v_/F_m_ values (0.618±0.002) than those from control conditions (Figs. 2C and D). Additional statistical information is included in supplemental materials (Table. S2).

**Fig. 2.**
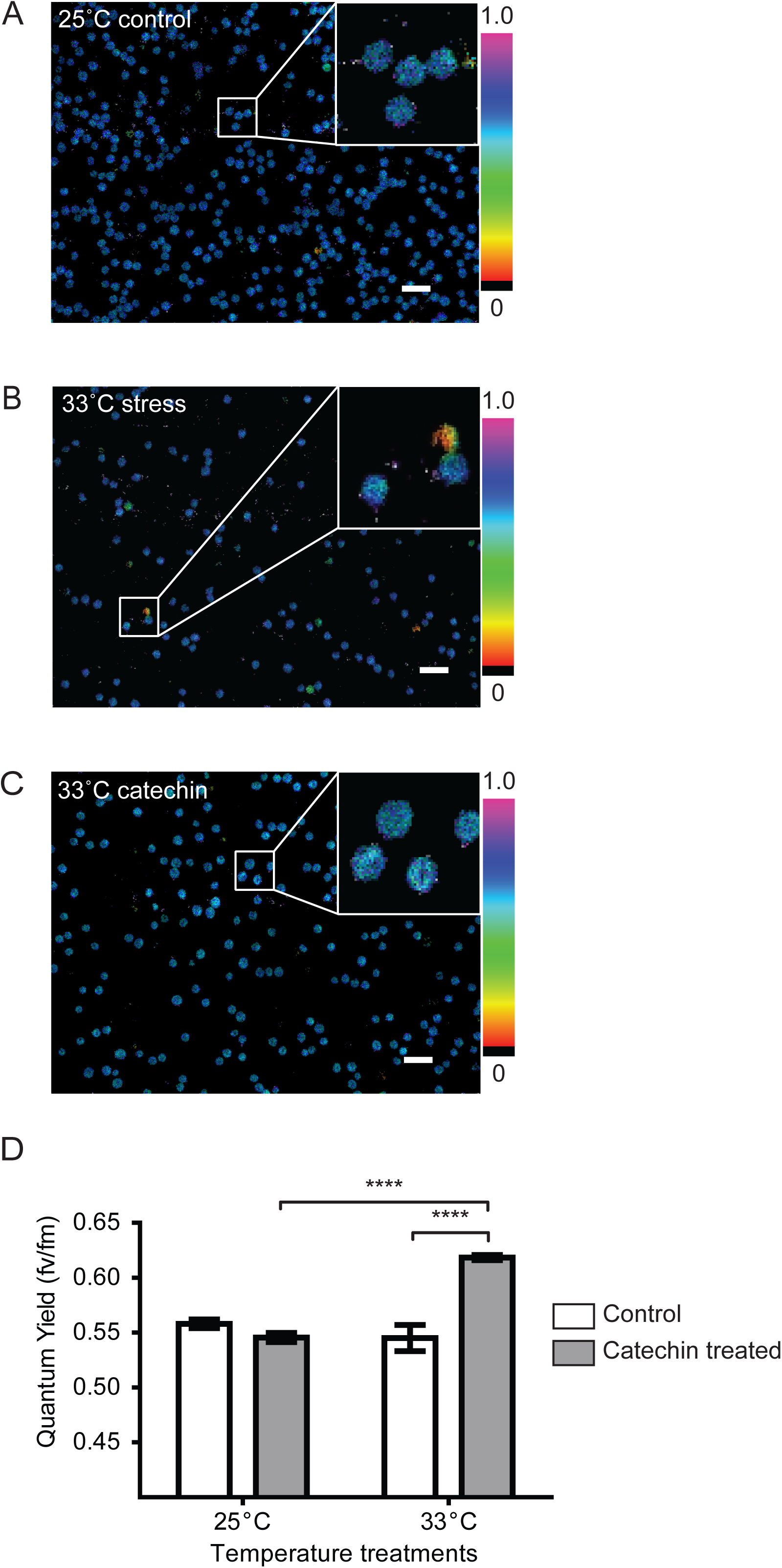
Maximum dark-adapted maximum quantum yield of photosystem II in *A. pallida* cell symbiosome. Representative PAM images of individual symbionts from (A) control, (B) thermal stress, (C) catechin + thermal stress treated anemones; associated colored gradient bars represent Fv/Fm value range (0.0 - 1.0). (D) Catechin + thermal stress treatment maximum photosynthetic quantum yield was significantly higher than their catechin control and thermal stress counterparts (2-Way ANOVA: temperature factor *P* < 0.0001; antioxidant factor *P* < 0.0001; temperature x antioxidant factor *P* < 0.0001). Bars represent means ± SE. *n* = 3 per treatment; sample size per *A. pallida* individual = 100 cells. Asterisks denote statistical significance of the post-hoc analyses. Scale bar = 20 μm.

### Natural antioxidant and ROS in scleractinian coral *P. astreoides*

Freshly isolated *P. astreoides* control symbionts incubated with the ROS sensitive dye had low (0.531±0.002 AU μm^−2^) fluorescence signaling when analyzed by confocal microscopy (Figs. 3A and B). Thermally stressed corals presented an increase in fluorescence signal (0.804±0.004 AU μm^−2^) in their symbionts (Figs. 3C and D). Corals incubated with catechin presented significantly reduced ROS levels (0.386±0.03 AU μm^−2^), similar to those observed in controls (Figs. 3E and F). Statistical analysis of multiple individual cells from several corals revealed that incubation with catechin reduced ROS levels in thermally stressed corals below that of control corals (Fig. 3G). Additional statistical information is included in supplemental materials (Table S3).

**Fig. 3.**
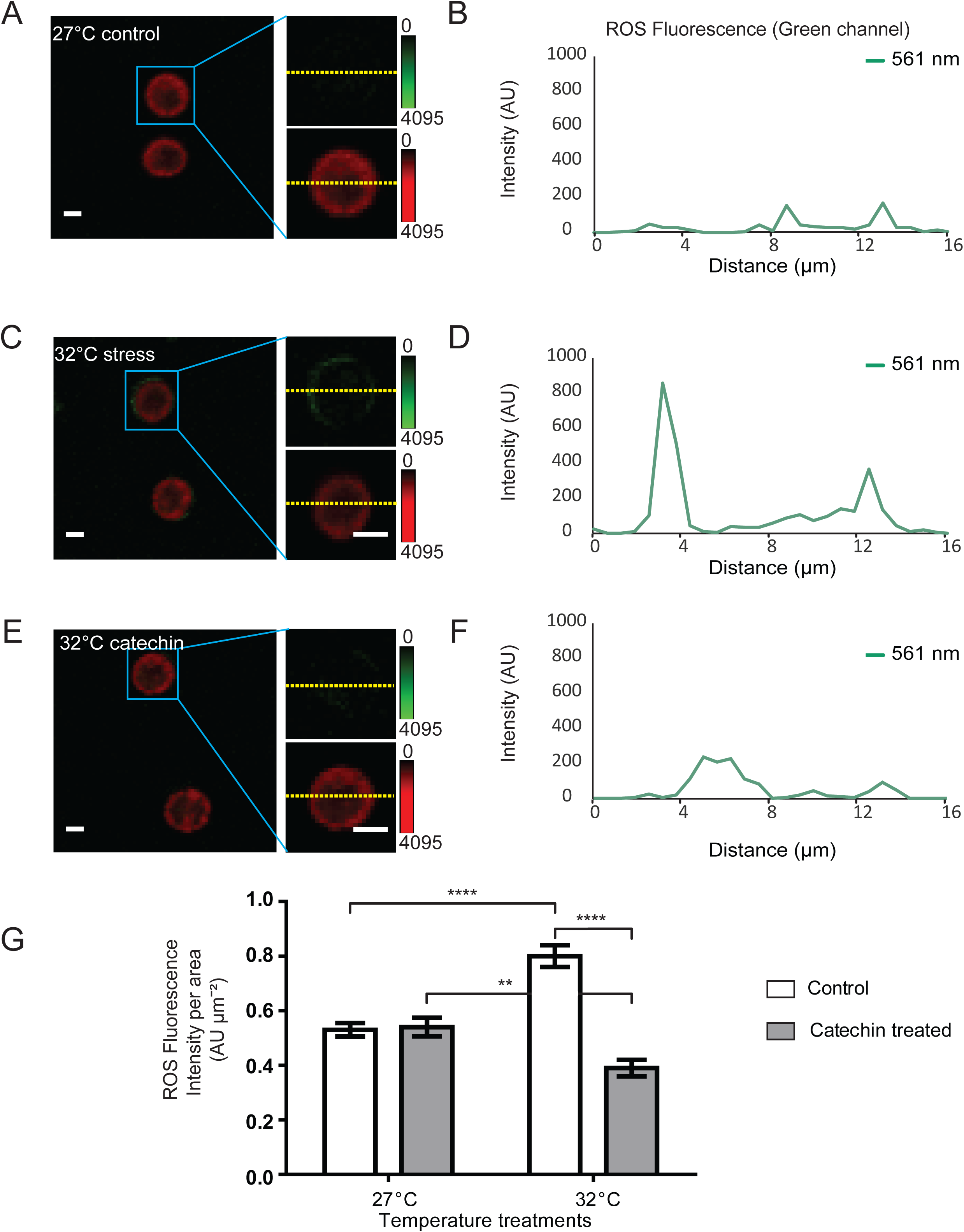
ROS levels in *P. astreoides* cell symbiosomes. (A) Control symbiosomes depicted low ROS fluorescence in the green channel (insert, top) and characteristic chlorophyll autofluorescence (insert bottom). (B) Line intensity profile from (A). (C) Thermal stressed symbionts emitted higher ROS fluorescence (top, insert). (D) Linear profile analysis from (C). (E) Catechin preincubations reduced thermal stress induced fluorescence similar to controls levels. (F) Line intensity profile from ROS insert in (E) showing a low ROS signal. (G) Catechin preincubation resulted a significant decrease in ROS fluorescence from thermal stress symbionts. (2-Way ANOVA: temperature factor *P*= 0.0861; antioxidant factor *P* < 0.0001; temperature x antioxidant factor *P* < 0.0001). Bars represent means ± SE. *n* = 4 per treatment; sample size per *P. astreoides* colony = 127-184 cells. Asterisks denote statistical significance of the post-hoc analyses. Scale bar = 5 μm.

### Dark-adapted maximum quantum yield of photosystem II of *P. astreoides*

Isolated *P. astreoides* symbiont F_v_/F_m_ values at 27 °C were 0.526±0.005 (Figs. 4A and D). This value was in agreement to previously described measurements in healthy corals subject to similar conditions (Warner *et al.*, 2006). Corals under thermal stress (32 °C for 4 days) however, presented decreased F_v_/F_m_ values to 0.429±0.007 (Figs. 4B and D). This result is also similar to data previously reported in stressed corals (Fujise *et al.*, 2014). Interestingly, F_v_/F_m_ values from thermally stressed corals in the presence of catechin were indistinguishable from control measurements (0.536±0.004), suggesting that the presence of catechin was sufficient to maintain basal F_v_/F_m_ values (Figs. 4C and D). Additional statistical information is included in supplemental materials (Table S4).

**Fig. 4.**
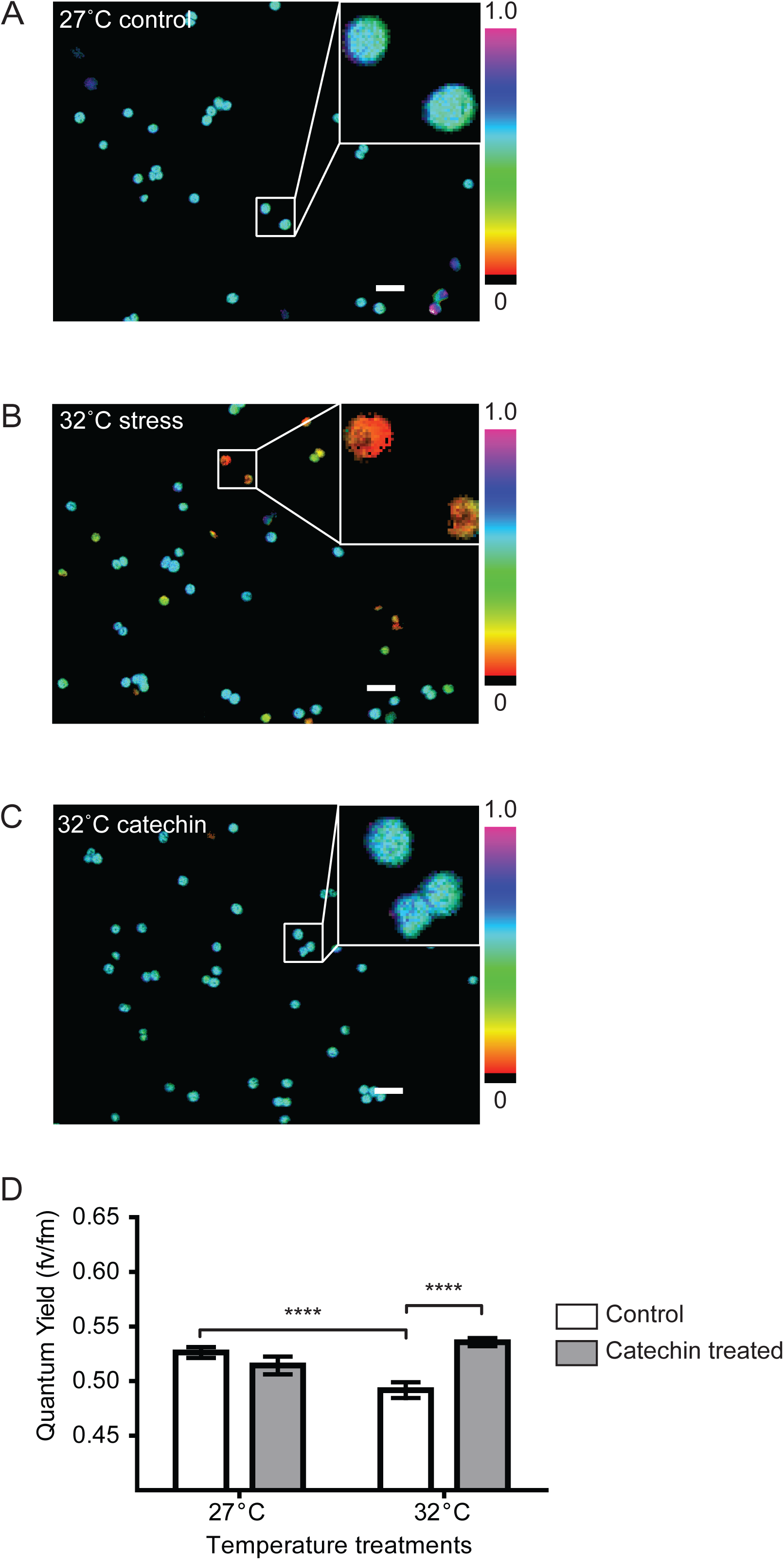
Dark-adapted maximum quantum yield of photosystem II in *P. astreoides* cell symbiosome. Representative fluorescence images obtained from PAM of (A) control, (B) thermal stress and (C) catechin + thermal stress cells. Colored gradient bars represent Fv/Fm range (0.0 - 1.0). (D) Stress treatment coral symbionts showed a statistically significant lower quantum yield than those in control and catechin + thermal stress treatments (2-Way ANOVA: temperature factor *P* = 0.3011; antioxidant factor *P* = 0.0122; temperature x antioxidant factor *P* < 0.0001). Bars represent means ± SE. *n* = 4 for (A) control and (C) 5 μM catechin + thermal stress; *n* = 3 for (D) catechin control treatments; *n* = 12 for (B) Stress treatment. Sample size per *P. astreoides* colony = 163-213 cells. Asterisks denote statistical significance of the post-hoc analyses. Scale bar = 20 μm.

### Symbiont cell density in *A. pallida* and *P. astreoides*

Next, we analyzed cell density before and after treatments. Thermally stressed sea anemones presented a significantly lower number of symbionts (3.713±0.04 × 10^7^ cells g^−1^) when compared to controls (1.085±0.15 × 10^8^ cells g^−1^) (Fig 5A). Catechin exposure did not prevent symbiont loss (4.305±0.06 × 10^7^ cells g^−1^) (Fig. 5A). In the case of *P. astreoides*, control samples exposed to catechin showed a decrease in total symbiont numbers (5.244±0.065 × 10^5^ cells ml^−1^). However, the presence of catechin significantly prevented further symbiont loss (4.913±0.046 × 10^5^ cells ml^−1^) when compared to thermally stressed samples (3.496±0.444 × 10^5^ cells ml^−1^), suggesting a palliative effect of catechin (Fig. 5B). Additional statistical information is included in supplemental materials (Tables S5 & S6).

**Fig. 5.**
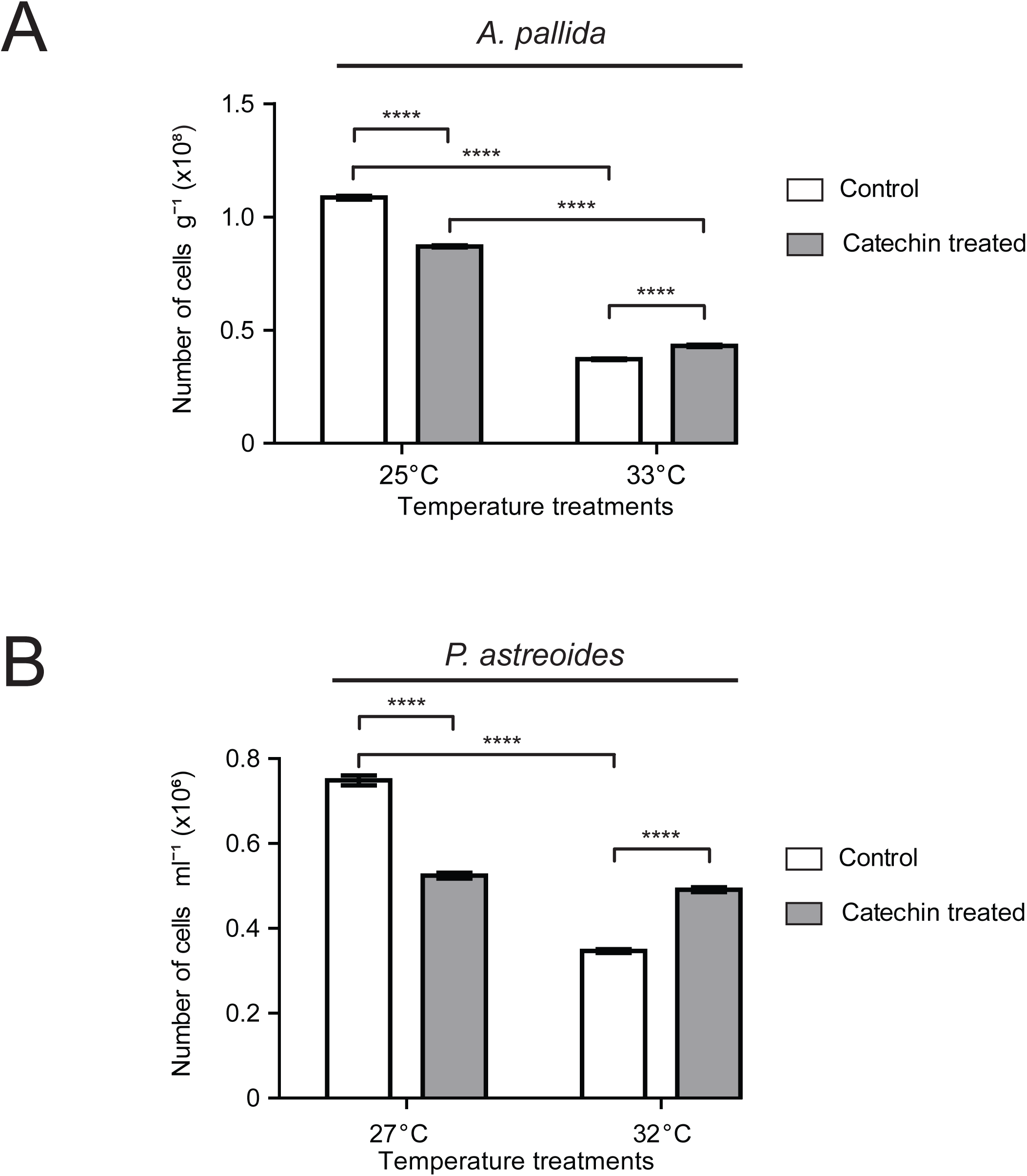
*A. pallida* and *P. astreoides* symbiont density. (A) Symbiont density from *A. pallida*. Bars represent means ± SE. *n* = 6-12. Sample size 1,224 – 3,606 cells. (2-Way ANOVA: temperature factor *P* < 0.0001; antioxidant factor *P* < 0.0001; temperature x antioxidant factor *P* < 0.0001). (B) Catechin prevented symbiont loss at high temperatures in *P. astreoides* (2-Way ANOVA: temperature factor *P* < 0.0001; antioxidant factor *P* < 0.0001; temperature x antioxidant factor *P* < 0.0001). Bars represent means ± SE. Asterisks denote statistical significance of the post-hoc analyses. *n* = 4 per treatment. Sample size 555-1198 cells.

### *A. pallida* symbiont relative frequency distribution

To further characterize the response of individual symbionts, we plotted the frequency distribution of cells based on their photosynthetic quantum yield. *A. pallida* controls had∼70% of symbionts in the range of 0.65 F_v_/F_m_, with a skewness of −5,2 (Fig. 6A); which was similar to thermal stress in the presence of catechin with 83% in the 0.65 bin, skewness −1.3 (Fig. 6B). For catechin controls, 83% cells were in the range of 0.55 F_v_/F_m_, with the remaining 17% in the 0.35-0.45 F_v_/F_m_ range and skewness of −1.5 (Fig. 6C). Thermally stressed symbionts yielded results similar to that of our stress plus catechin treatment with the 68% of the cells in the 0.55 F_v_/F_m_ bin and a skewness of −3.8 (Fig. 6D).

**Fig. 6.**
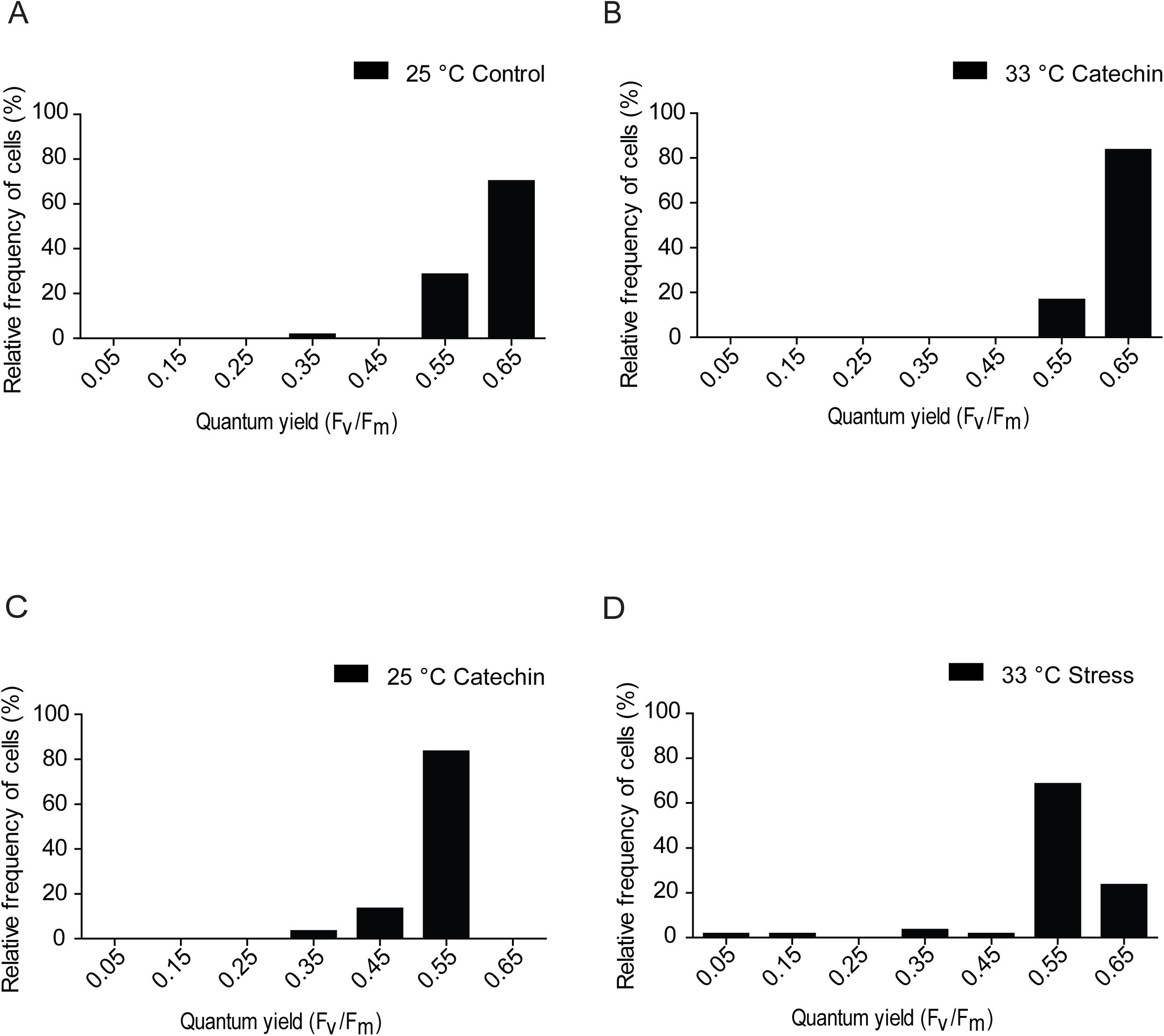
*A. pallida* symbiont photosynthetic quantum yield based distribution. Single cell quantum yield data was plotted relative total frequency (%) (A) Control treatment. (B) Thermal stress + catechin treatment s (C) Catechin control treatment. (D) Thernal stress treatment Bin width = 0.1.

### *P. astreoides* symbiont relative frequency distribution

*P. astreoides* controls exhibited ∼88% of the symbionts at the 0.55 F_v_/F_m_ bin with a skewness of −5.59 (Fig. 7A). Stressed plus catechin treatment symbionts behaved similarly to those observed in the control treatment where ∼89% were also at 0.55 F_v_/F_m_ and similar skewness of −6.15 (Fig. 7B). The catechin control, on the other hand, had several groups ranging from 0.15 – 0.45 and a main group at 0.65 F_v_/F_m_ and skewness of −1.6 (∼80%; Fig. 7C). Lastly, stress treatment also had similar distribution with 68% of the cells at 0.55 F_v_/F_m_ and a skewness of −2.32 (Fig. 7D).

**Fig. 7.**
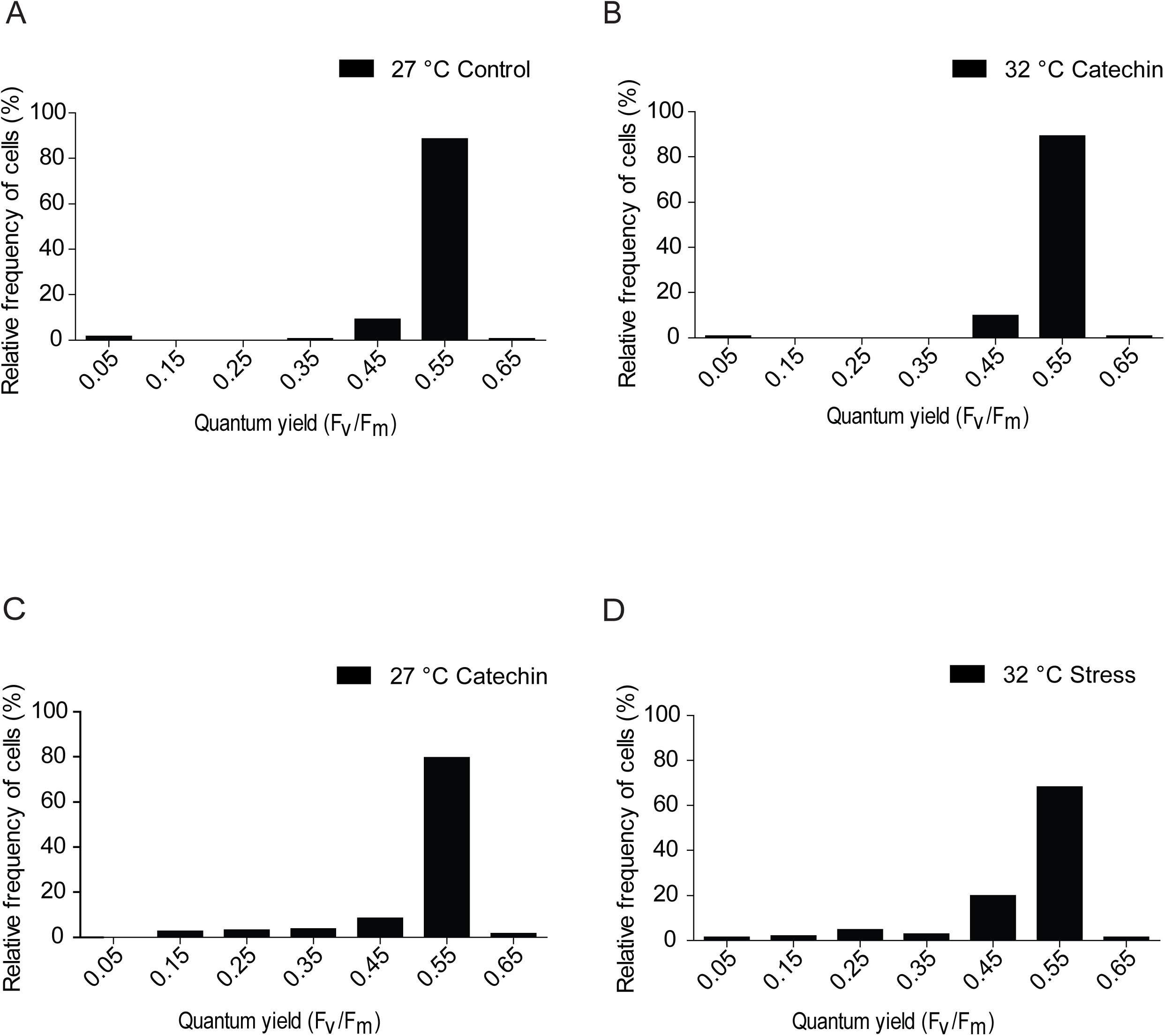
*P. astreoides* symbiont photosynthetic quantum yield based distribution. Single cell quantum yield data was plotted relative to total frequency (%) (A) Control treatment. (B) catechin + Thermal stress treatment (C) Catechin control treatment. (D) Thermal stress treatment Bin width = 0.1.

## DISCUSSION

To further understand the molecular events leading to symbiont exoulsion, we tested the hypothesis that thermally-induced increases in ROS are sufficient to damage the photosynthetic machinery of the symbionts and induce symbiont loss. Photosynthetic degradation, as measured by quantum yield of the photosystem II (F_v_/F_m_) and symbiont density were analyzed in the model system *A. pallida* and the coral *P astreoides*. We also tested the hypothesis that antioxidants could be used to prevent these events. Here we show that thermal stress increased ROS production in both anemones and corals, and that this increase was accompanied by a reduction in the number of symbionts present in the host tissue. Most importantly, we found that treatment with a naturally-occurring antioxidant can reduce ROS in coral tissues and possibly protect the symbiont loss.

Previously, studies have suggested ROS as a possible molecular mechanism triggering coral bleaching, and their reduction can have a positive effect in some species at the macroscopic level (Lesser, 1997). Our experimental results complement these studies by revealing that thermally-induced increases in ROS resulted in a lowered F_v_/F_m_ in symbionts from *A. pallida* and *P. astreoides.* Our single cell approach was not designed to measure ROS concentrations; however, the increases in ROS observed when exposed to thermal stress were similar in magnitude and kinetics in these two species. Interestingly, a single dose of the naturally occurring antioxidant catechin, was sufficient to reduce ROS levels in host cells while preventing symbiont F_v_/F_m_ degradation in both *A. pallida* and *P astreoides*. Furthermore, catechin also mitigated symbionts loss, but only from thermally stressed *P. astreoides* and not from *A. pallida*. It is interesting to note that catechin reduced the ROS signal in control temperature samples below untreated samples, indicating background basal levels of ROS or mild stress in these samples. However, this also resulted in loss of symbionts, suggesting a possible physiological role of these molecules during basal events such as homeostatic and immune signaling responses (Villanueva and Kross, 2012). We also saw a loss of symbionts under control conditions when we exposed our organisms to catechin in the absence of a temperature increase, whereas exposure to both thermal stress and catechin had results similar to control conditions in both *A. pallida* and *P. astreoides*. This suggests that catechin by itself could be damaging to photosynthetic pathways, perhaps by reducing ROS below a functional level required for signaling or other cellular processes, which in turn could trigger the loss of symbionts.

Additionally we found that the photosynthetic quantum yield distribution of *A. pallida* symbiont cells had a decline in cells while being exposed to catechin in control temperatures. This result further shows that catechin, by itself may be damaging to the QY. In the case of *P. astreoides,* the photosynthetic quantum yield distribution of symbiont cells under thermal stress was similar to that of our control temperature plus catechin treatment; while the thermal stress treatment was similar to our control. Being exposed to catechin without thermal stress may be as damaging to the photosynthetic quantum yield of cells as if they were exposed to thermal stress.

From the molecular perspective, our data suggest that high levels of ROS are sufficient to elicit bleaching, while a reduction in their levels can mitigate thermally induced bleaching in the coral *P. astreoides*. Can antioxidants be utilized to prevent coral bleaching in their natural environment? Such an appealing idea will need careful consideration and further investigation in other coral and animal species before any field testing. For example, we showed that different species can respond differently to antioxidant treatments. Why did catechin prevent bleaching in *P. astreoides* and not in *A. pallida*? This indicates that there may be underlying differences in the bleaching responses or mechanisms between the anemone *A. pallida* and hard corals and therefore using *A. pallida* as a model system to investigate coral bleaching may be misleading. Processes such as DNA damage and apoptosis have been described during coral bleaching, underlying the complexity of the interplay between host and symbionts during stress (Lesser, 2011). Future experiments expanding our understanding of coral bleaching and the role of oxidative stress at the cellular and molecular level with different coral species should help us answer these important questions.

## Acknowledgments

We thank Neidibel Martinez for technical assistance in the laboratory, Dr. Amelia Merced and Carlos Vázquez for microscopy assistance, personnel at Marine Biological Laboratory’s Marine Resources Center for equipment and assistance, and members of Loretta Roberson’s laboratory for maintenance and care of tanks. We thank Dr. Carlos Garcia (Clemson University) for his intellectual feedback and comments on our research.

This research was supported by a National Science Foundation grants HRD #1137725 and DBI #1337284. GAY is supported by grants from NIH DA037924 and DA040920.

---

Supplemental Fig. 1. ***Symbiodinium* sp. autofluorescence spectrum.** Isolated coral symbionts emission spectrum analyzed by confocal microscopy shows an autofluorescence peak in the 670-690 nm region (Red zone) and relatively low autofluorescence in the 565-nm CellROX ^®^ Orange region (Orange zone). *n* = 1; sample size = 4 cells.

